# Preclinical Screening of Splice-Switching Antisense Oligonucleotides in PDAC Organoids

**DOI:** 10.1101/2023.03.31.535161

**Authors:** Ledong Wan, Alexander J. Kral, Dillon Voss, Adrian R. Krainer

**Affiliations:** Cold Spring Harbor Laboratory, Cold Spring Harbor, NY 11724, USA; Stony Brook University, Stony Brook, New York

## Abstract

Aberrant alternative splicing is emerging as a cancer hallmark and a potential therapeutic target. It is the result of dysregulated splicing factors or genetic alterations in splicing-regulatory *cis*-elements. Targeting individual altered splicing events associated with cancer-cell dependencies is a potential therapeutic strategy, but several technical limitations need to be addressed. Patient-derived organoids (PDOs) are a promising platform to recapitulate key aspects of disease states and to facilitate drug development for precision medicine. Here, we report an efficient antisense-oligonucleotide (ASO) transfection method to systematically evaluate and screen individual splicing events as therapeutic targets in pancreatic ductal adenocarcinoma (PDAC) organoids. This optimized delivery method allows fast and efficient screening of ASOs that reverse oncogenic alternative splicing. In combination with advancements in chemical modifications and ASO-delivery strategies, this method has the potential to accelerate the discovery of anti-tumor ASO drugs that target pathological alternative splicing.

## Introduction

Pre-mRNA splicing is catalyzed in the cell nucleus by the spliceosome, a large and dynamic ribonucleoprotein complex that precisely excises the introns and joins the exons to form mature mRNAs. High-throughput RNA sequencing studies showed that the pre-mRNAs from 90–95% of human genes undergo alternative splicing, prominently contributing to transcriptomic and proteomic diversity (1, 2). Coordinated alternative splicing confers essential physiological functions that are critical for developmental processes, tissue homeostasis, and cellular identity.

Aberrant alternative splicing is associated with various diseases, notably cancer, and is attributable to dysregulated splicing factors or to mutations in splicing-regulatory *cis*-elements (3). Cancer cells often exhibit aberrant alternative-splicing profiles, leading to the production of isoforms that increase cell proliferation, migration, or resistance to apoptosis, or to the loss of isoforms that block these processes (2). Increased expression of several RNA-binding proteins involved in splicing is a distinguishing feature for the most aggressive subtype of pancreatic cancer, in conjunction with mutations in *KRAS* and *TP53* (4). Several potential therapeutic strategies—including targeting individual dysregulated splicing factors or the entire spliceosomal machinery, as well as targeting individual altered splicing events—have been proposed, either alone or in combination with immunotherapy (2, 5, 6). However, several obstacles need to be addressed: i) targeting individual splicing factors involved in spliceosome assembly will affect the transcriptome globally, with likely off-target toxicity in normal tissues; ii) targeting individual alternative splicing events is currently hampered by insufficient knowledge about pivotal alternative-splicing events in cancer.

Although genetically engineered mouse models have greatly contributed to the understanding of cancer development, some biological processes cannot be fully recapitulated and correctly interpreted using mouse models. Patient-derived organoids (PDOs), a stem-cell-derived 3D culture system, recapitulate the architecture and physiology of human healthy or diseased organs, thus helping to overcome the limitations of animal models in addressing specific aspects of human biology and disease (7). Additionally, these organoid models can be used as a platform for accurately assessing precision-medicine-based drug development (8). PDOs offer numerous advantages, including preservation of patient-to-patient variability, ease of genetic manipulation, xenotransplantation, targeted development of personalized drugs, and matched drug sensitivity between *in vitro* culture and *in vivo* xenografts. Indeed, several studies have recently leveraged PDOs for high-throughput drug screening (8–10).

Antisense oligonucleotides (ASOs) are synthetic, chemically modified, short chains of nucleotides with the potential to regulate gene expression through several mechanisms, such as steric blocking resulting in splice-switching or translation inhibition, and RNase-H-mediated mRNA cleavage (11). We and others have developed splice-switching ASOs that have been successfully used in the clinic, e.g., nusinersen for spinal muscular atrophy (SMA) (12), and eteplirsen, golodirsen, viltolarsen, and casimersen for Duchenne muscular dystrophy (DMD) (13, 14). Unlike SMA and DMD, which are monogenic diseases that can be treated by targeting a particular exon, cancers usually display hundreds of abnormal alternative splicing events, some of which may be pro-oncogenic, whereas the rest may be passenger changes. Validating and characterizing the functional consequences of these potential oncogenic events in clinically relevant models is critical to drive innovative therapeutic modalities.

Here, we report a versatile transfection-based method that can be applied for systematic evaluation and screening of ASO candidates and their splicing targets in human PDAC organoid model systems. ASO free uptake is frequently hindered by very slow uptake kinetics, the high ASO concentrations required, and cell-type variability, making it impractical for initial ASO-screening purposes. Our method optimizes the uptake efficiency of ASOs to facilitate effective preclinical evaluation of their therapeutic potency, while preserving the physiologically relevant organoid architecture. This delivery method allows fast and efficient screening of ASO candidates against oncogenic alternative-splicing targets.

In combination with advances in oligonucleotide chemical modifications, conjugation, and other *in vivo* ASO-delivery strategies, we suggest that our method will help accelerate the discovery of anti-tumor ASO drugs more broadly, beyond targeting pathological alternative splicing.

## Results

### Baseline free-uptake efficiency of ASOs in a 2D cell line

To standardize our delivery pipeline, we used two splice-switching ASO sequences that our lab has extensively characterized for targeting *SMN2* and *PKM* pre-mRNA, respectively (12, 15–17). In this study we used commercially obtained ASOs, uniformly modified with 2’-*O*-methoxyethyl (MOE) ribose, phosphorothioate backbone, and 5-methyl cytosines, which is the same set of modifications present in nusinersen, the first FDA-approved drug for the treatment of SMA (12).

We used an *SMN2* pre-mRNA splice-switching ASO (ASO-S) as an example of an ASO with high splice-switching potency. ASO-S is 2-nt longer than nusinersen and has very similar activity (15). These ASOs target the *SMN2* pre-mRNA, which undergoes alternative splicing. The predominant spliced mRNA isoform lacks exon 7, which is the last coding exon. Translation of this mRNA incorporates a peptide from exon 8 that forms a degron at the SMN C-terminus, causing protein degradation (Figure 1A, (18)). ASO-S binds to *SMN2* intron 7, blocking binding of a splicing repressor and promoting efficient exon 7 inclusion in the spliced *SMN2* mRNA, which is translated into the full-length SMN protein (Figure 1A, (12)).

**Figure 1:**
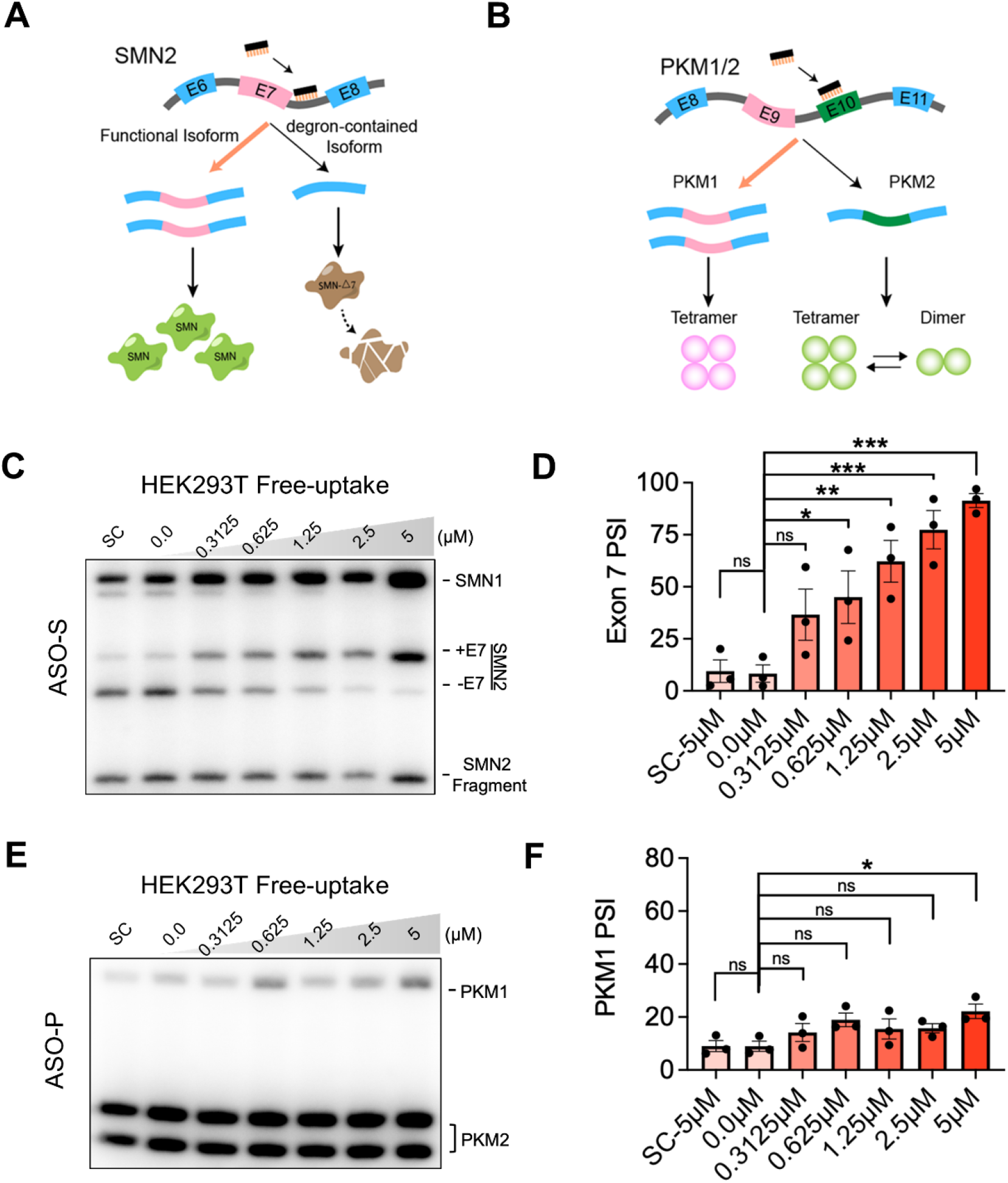
Lead ASOs exhibit different splice-switching efficiency in HEK293T cells. **(A)** Schematic of ASO-mediated splice-switching in *SMN2*. **(B)** Schematic of ASO-mediated splice-switching in *PKM*. **(C)** Radioactive RT-PCR of *SMN1/2* mRNA in HEK293T cells treated by free uptake of ASO-S. **(D)** PSI quantification of **(C)** (n=3). **(E)** Radioactive RT-PCR of *PKM* mRNA in HEK293T cells treated by free uptake of ASO-P. **(F)** PSI quantification of **(E)** (n=3). SC, scrambled control. One-way ANOVA, \**P*<0.05, \*\**P*<0.01, \*\*\**P*<0.001. Error bars represent ± SEM.

In addition to ASO-S, we also used ASO-P (*PKM2*-targeting ASO) as another splice-switching ASO currently undergoing pre-clinical development, and one that displays comparatively modest splice-switching effects for its target, relative to that of ASO-S. The *PKM* gene, which comprises 12 exons, encodes two pyruvate kinase M isoforms (PKM1 and PKM2) through mutually exclusive alternative splicing of exons 9 and 10 (Figure 1B, (16)). PKM1 is expressed in terminally differentiated, non-proliferating cells, whereas PKM2 is highly expressed in proliferating embryonic cells. In many types of cancer, expression is shifted from PKM1 to PKM2, resulting in an increase in aerobic glycolysis, also known as the Warburg effect (19). Selectively targeting *PKM* in cancer cells is an attractive therapeutic modality. ASO-P downregulates PKM2, while simultaneously upregulating PKM1, resulting in apoptosis in glioblastoma cells and cell-cycle arrest in hepatocellular carcinoma (HCC) cells (16, 20). Given that ASO-P’s free-uptake and splice-switching efficiencies are suboptimal, we sought to optimize and standardize a method for delivering ASOs targeting different alternative-splicing events, which would allow us to evaluate the functional significance and therapeutic relevance of these events.

We first examined the efficiency of ASO-S and ASO-P in HEK293T cells by free uptake. Cells were incubated with varying ASO concentrations, ranging from 0 to 5 μM, for 3 days. Then, the ASO and medium were replenished, and the cells were incubated for an additional 2 days, prior to RNA extraction and analysis by radioactive RT-PCR. To examine the splicing changes in *SMN2*, cDNAs were digested with DdeI to separate the *SMN2* and *SMN1* PCR products (12). After digestion and separation by 5% native PAGE, bands of 398 bp and 344 bp are observed that correspond to included or skipped *SMN2* exon 7, respectively (Figure 1C). Consistent with our previous studies, ASO-S promoted *SMN2* exon 7 inclusion in a dose-dependent manner, with 5 μM ASO-S resulting in around 90% full-length *SMN2* isoform expression (Figure 1D).

To analyze *PKM* splicing, primers annealing to the flanking constitutive exons 8 and 11 were used to reverse-transcribe and amplify both *PKM1* and *PKM2* isoforms. After amplification, the 398 bp PCR products were digested with PstI to distinguish the *PKM1* and *PKM2* isoforms (16). Two PCR fragments of 213 bp and 185 bp are generated for *PKM2*, whereas *PKM1* remains intact. ASO-P elicited modest splice-switching of *PKM* only at the highest concentration tested (Figure 1E, 1F), whereas the above ASO-S showed marked splice-switching starting at 0.3125 μM. These results establish the baseline efficiency of these ASOs in 2D cell culture, which is not affected by variables associated with the use of Matrigel and the 3D architecture of organoids.

### Limited splice-switching efficiency of ASOs in PDAC organoids

We next sought to determine the effectiveness of ASO delivery in human PDAC organoids. We first tested ASO delivery via free uptake, due to its ease and relevance for clinical applications (20, 21). We used hF23 and hT60 organoids, corresponding to classical and basal molecular PDAC subtypes, respectively (22). Organoids were seeded in Matrigel, and culture medium was added after the matrix solidified. ASO-containing culture medium was replenished on Day 3. After 5 days of incubation, the organoids were harvested (Figure 2A).

**Figure 2:**
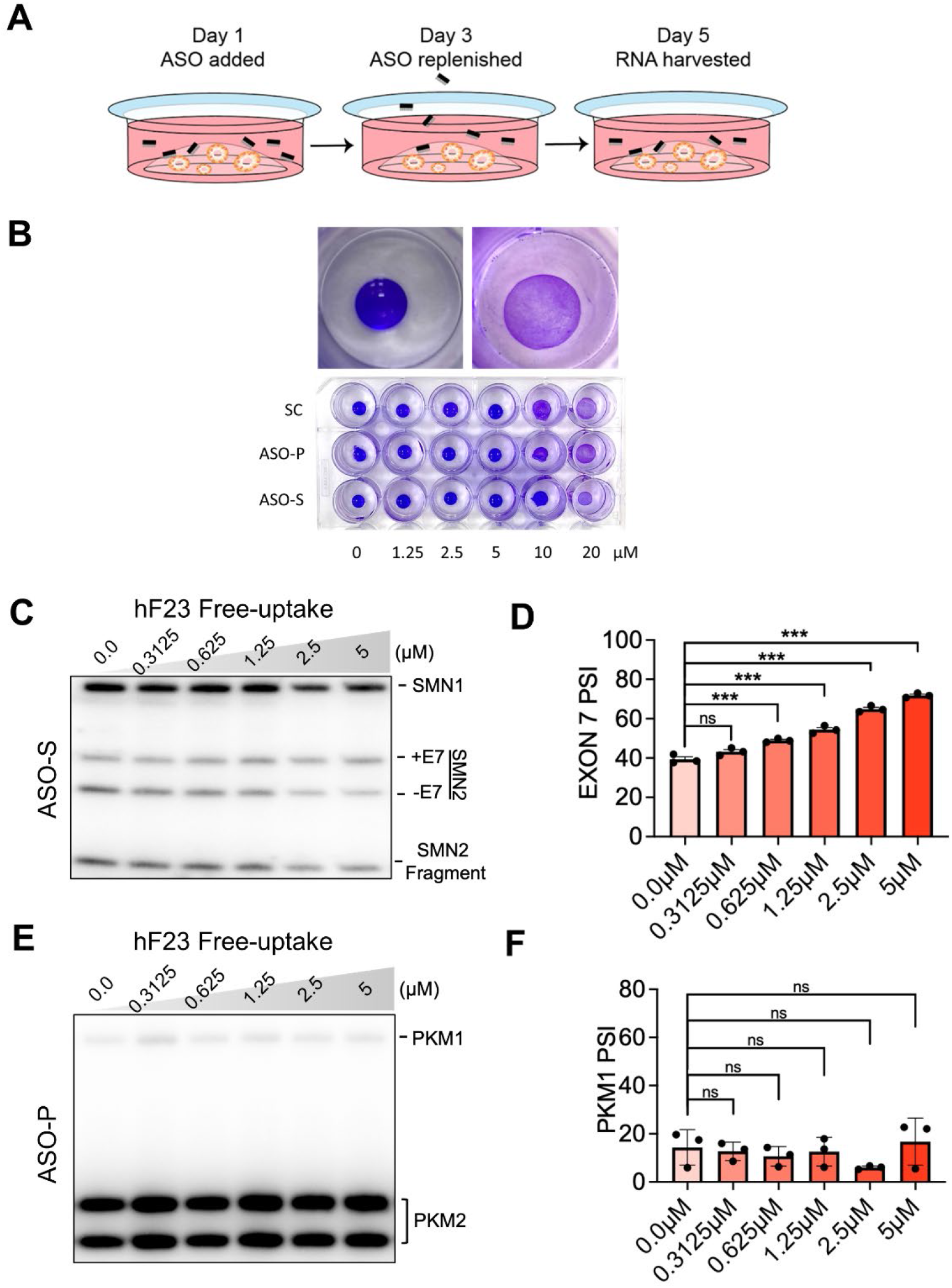
Limitations of ASO free uptake in PDAC organoids. **(A)** Schematic of ASO free uptake in organoid culture. **(B)** Matrigel domes were treated with different concentrations of scramble control, ASO-P and ASO-S ASOs for 5 days. **(C)** Radioactive RT-PCR of *SMN1/2* mRNA in hF23 organoids treated by free uptake of ASO-S. **(D)** PSI quantification of **(C)** (n=3). **(E)** Radioactive RT-PCR of *PKM* mRNA in hF23 organoids treated by free uptake of ASO-P. **(F)** PSI quantification of **(E)** (n=3). One-way ANOVA, \*\*\**P*<0.0001. Error bars represent ± SEM.

During the incubation, we noticed the collapse of the Matrigel domes at high ASO concentrations. Even in the absence of organoids, incubation with concentrated ASO resulted in loose or dissolved Matrigel, as shown by crystal-violet staining (Figure 2B). Therefore, the practical ASO concentration in free-uptake experiments is limited by the Matrigel properties, with a loss of organoid architecture at high ASO concentrations.

The *SMN2* gene was expressed in hF23 but not in hT60 organoids, reflecting the absence of this non-essential paralog in some individuals (23). We found that free uptake of ASO-S in organoids resulted in slight splicing changes in *SMN2*. Exon 7 inclusion increased by ~25% at 5 μM ASO in hF23 organoids, whereas it increased by >80% in HEK293T cells (Figure 2C, 2D). In addition, *PKM1/2* splicing was minimally affected by ASO-P free uptake in hF23 and hT60 organoids (Figure 2E, 2F, S2A, S2B). We conclude that organoid cultures are limited in their ability to withstand high ASO concentrations without losing important 3D features, which results in inefficient ASO uptake and/or effectiveness. We therefore sought to establish a more efficient delivery method that preserves organoid 3D architecture and requires less time and lower ASO concentrations.

### ASO free uptake in Matrigel-diluted PDAC organoid cultures

High-throughput drug screening using PDAC organoids often employs a hybrid 2D/3D plating strategy involving a Matrigel bed and a diluted mixture of Matrigel with organoid culture medium (22, 24). We first tested whether using this strategy could promote ASO uptake. We dissociated human PDAC organoids into single cells and resuspended them with 10% Matrigel/human complete feeding medium and ASO at varying concentrations. Organoid/Matrigel/media/ASO mixtures were then plated in 96-well plates on top of a 50% Matrigel/DPBS bed. After incubation for 5 days, organoids were harvested and the extracted RNA was analyzed by radioactive RT-PCR (Figure 3A). We observed a slight increase in *SMN2* exon 7 inclusion after treatment with ASO-S in the hF23 organoid line. However, the increase was not statistically significant, which we attribute to the large variance between samples. The efficiency of ASO-S in the Matrigel-diluted culture was slightly less than by free uptake (Figure 3B and 3C). In addition, ASO-P failed to elicit significant splice-switching of *PKM* (Figure 3D, 3E, S2C, S2D), further highlighting the need for a standardized method for efficiently delivering ASOs into organoids, while preserving their functional and clinically relevant features.

**Figure 3:**
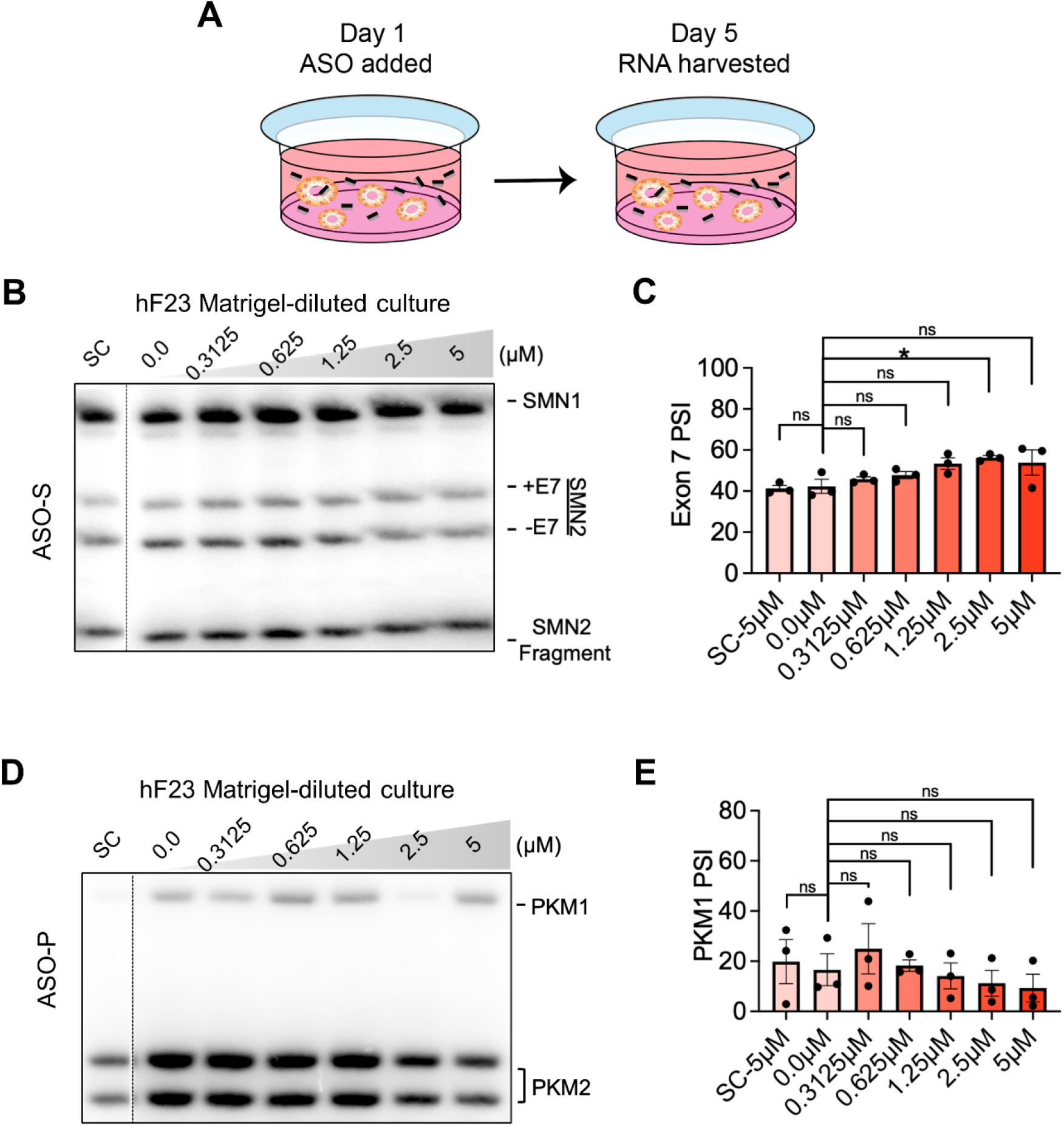
Splice-switching efficiency of ASO free uptake in Matrigel-diluted organoid culture. **(A)** Schematic of ASO free uptake in the Matrigel-diluted organoid culture. **(B)** Radioactive RT-PCR of *SMN1/2* mRNA in Matrigel-diluted cultured hF23 organoids treated by free uptake of ASO-S. **(C)** PSI quantification of **(B)** (n=3)**. (D)** Radioactive RT-PCR of *PKM* mRNA in Matrigel-diluted cultured hF23 organoids treated by free uptake of ASO-P. **(E)** PSI quantification of **(D)** (n=3). SC, scrambled control. One-way ANOVA, \**P*<0.05. Error bars represent ± SEM.

### Optimized transfection method reproducibly delivers ASOs in PDAC organoids

In general, transfection markedly improves ASO uptake and effectiveness in 2D-cultured cells. We found that transfection gave comparable splice-switching of *SMN2* and *PKM* pre-mRNAs by ASO-S and ASO-P, respectively, at equivalent low-dose levels in 2D cells. Both ASOs promoted ~50% changes in their targeted splicing events in HEK293T cells (Figure S3). In line with our previous reports (16), the extensive splice-switching between PKM1/2 isoforms was accompanied by the appearance of the PKMds (double-skipped) isoform lacking both exon 9 and exon 10 in the spliced mRNA (Figure S3C).

Given the high efficiency of ASO delivery by transfection, we sought to apply it in the PDAC organoid culture. To improve the transfection efficiency, we first dissociated the organoids into single-cell suspensions. The single cells were then spun down and resuspended in lipofectamine 3000-ASO mixture and incubated at 37 °C at 5% CO2 for 8 hours. The cells were then spun down and resuspended in Matrigel using standard organoid culture conditions (Figure 4A). After 72 h the re-formed organoids were harvested, and extracted RNA was analyzed by radioactive RT-PCR. We checked that our method did not compromise PDAC-organoid morphology or molecular features, by using bright-field imaging and performing RT-qPCR for ductal lineage markers (25). Our data demonstrate that transfection did not perturb the characteristic organoid features (Figure 4B and 4C).

**Figure 4:**
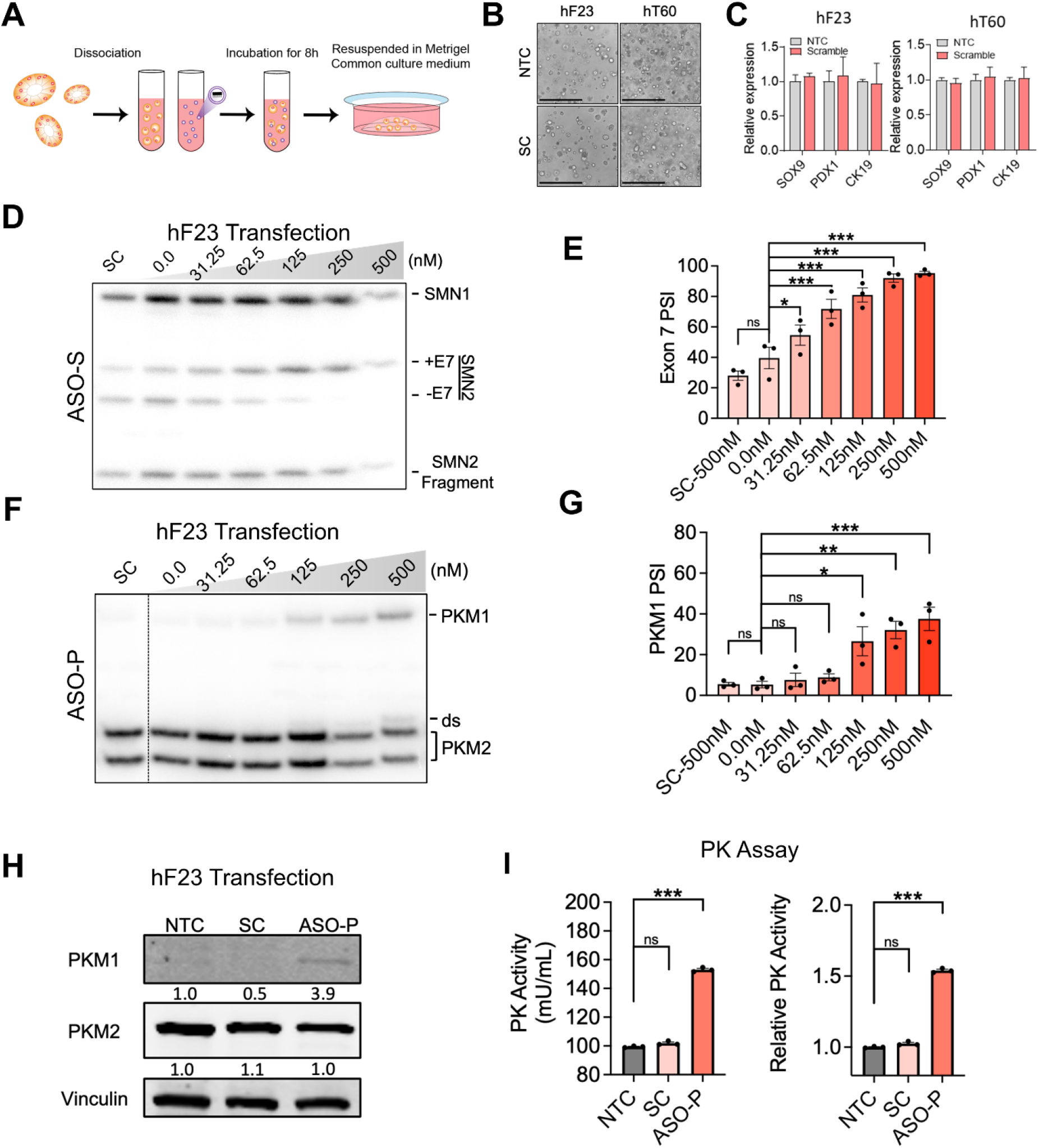
Transfection-based ASO delivery in PDAC organoids. **(A)** Schematic of transfection-based ASO delivery in PDAC organoids. **(B)** Bright-field image of PDAC organoids after ASO transfection. **(C)** qPCR of PDAC organoid markers in hF23 and hT60 transfected with SC ASO. **(D)** Radioactive RT-PCR of *SMN1/2* mRNA in hF23 transfected with ASO-S. **(E)** PSI quantification of **(D)** (n=3)**. (F)** Radioactive RT-PCR of *PKM* mRNA in hF23 transfected with ASO-P. **(G)** PSI quantification of **(F)** (n=3)**. (H)** Western blot of PKM1 and PKM2 in hF23 transfected with 125 nM ASO. Fold change was calculated by normalizing band intensity to vinculin loading control (n=2). **(I)** Pyruvate Kinase (PK) activity and relative PK activity in hF23 transfected with 125 nM ASO-P. NTC, no-treatment control; SC, scrambled control. One-way ANOVA, \**P*<0.05, \*\**P*<0.01, \*\*\**P*<0.001. Error bars represent ± SEM.

The optimized transfection protocol in hF23 organoids markedly improved the *SMN2* splicing change induced by ASO-S, which was comparable to that of transfection in HEK293T cells (Figure 4D, 4E, S3A, S3C). Furthermore, transfection of ASO-P likewise induced significant splice-switching in *PKM* in a dose-dependent manner (Figure 4F, 4G, S4A, S4B) in both hF23 and hT60 organoid lines.

We also validated the corresponding PKM protein isoform changes in PDAC organoids transfected with ASO-P. hF23 and hT60 lines transfected with 125 nM ASO-P exhibited 4-8 fold elevated protein levels of PKM1 (Figure 4H and S4C). As expected—because PKM1 is more catalytically active than PKM2 (16)—we observed a robust increase in pyruvate kinase (PK) activity (Figure 4I and S4D).

In sum, we described a fast and simple transfection method to reproducibly elicit ASO-mediated splice-switching in target pre-mRNAs. This method should be applicable to other alternative splicing targets, and we expect that it will accelerate the screening of precision ASO therapeutics for PDAC.

## Discussion

High-throughput drug screening has been an attractive application of PDAC organoids, due to their ability to recapitulate the drug sensitivity of individual patients (26). Here, we established a method for cost-effectively delivering splice-switching ASOs to human PDAC organoids, which we hope will help to develop ASO therapeutics as next-generation cancer therapeutics (Figure 5). With its optimized efficiency, this platform makes it possible to interrogate physiologically and pathologically relevant alternative splicing events in PDAC. This in turn should facilitate the identification of critical therapeutic targets for clinical development. Given that PDAC organoids preserve clinically relevant features (7)—e.g., tumor heterogeneity and drug response—ASO screening in organoids can facilitate the validation of target AS events and the identification of lead ASOs for personalized-drug development.

**Figure 5:**
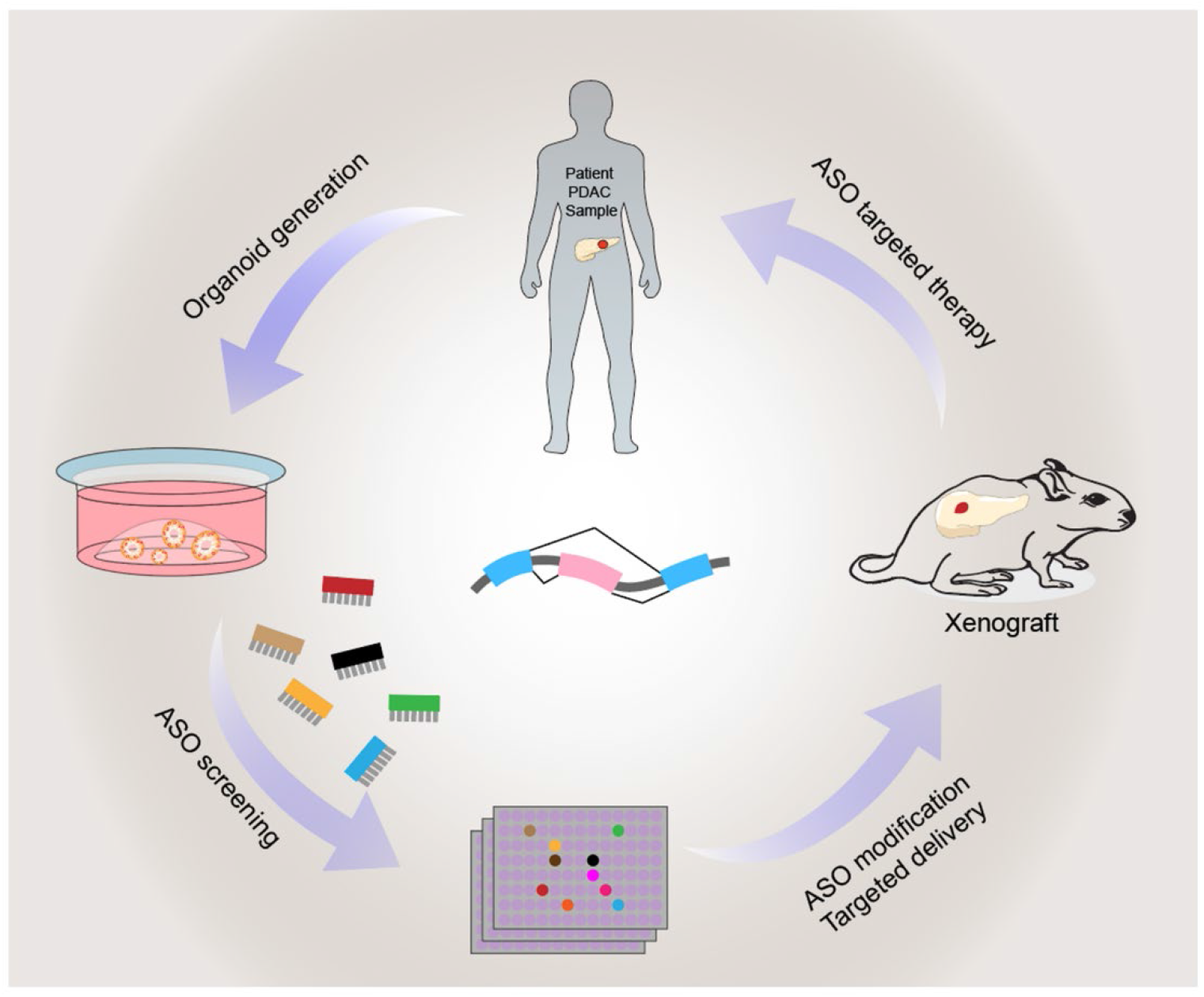
Schematic overview of the techniques integrated with the objective of improving personalized medicine for pancreatic cancer: organoid isolation culture and application for ASO drug screening. Tumor-derived organoids serve as a drug-screening platform to identify tumor-specific alternative splicing vulnerabilities.

The method described herein should be compatible with other advances in antisense technology. For example, conjugation with antibodies (27), GalNAc (28), or aptamers (29) might enhance ASO delivery in organoids. Lead ASOs identified using our method can be optimized by subsequent conjugation of various ligands to enhance delivery to target tissues. Additionally, ASOs have been developed for manipulating gene expression through multiple mechanisms, in addition to alternative splicing, such as activating RNase-H mediated knockdown, inhibiting nonsense-mediated mRNA decay, influencing polyadenylation to modify mRNA stability, or regulating the translation of target mRNAs (30–32). Our delivery method can readily be extended to evaluate these other modalities. Lastly, given the broad application of organoids for studying cancer, infectious diseases, and genetic disorders (7, 33, 34), we expect that our method for PDAC organoids can be adapted to explore ASO therapeutic potential in other disease contexts.

## Acknowledgments

We gratefully acknowledge Tuveson laboratory and the Organoids Shared Resources at the Cold Spring Harbor Laboratory (CSHL) for human PDAC organoid culture. Funding: NCI Program Project Grant CA13106 Project 2 to A.R.K.; CSHL Shared Resources used in this work were funded in part by NCI Cancer Center Support Grant 5P30CA045508.

## Author Contribution

Conceptualization: L.W., A.J.K., A.R.K.; Methodology: L.W., A.J.K., D.V., A.R.K.; Investigation: L.W., A.J.K., D.V.; Writing: L.W., A.J.K., A.R.K.; Supervision: A.R.K.

## Materials and Methods

### Tissue Culture

HEK293T cells were cultured in DMEM supplemented with 10% FBS and 1% Penicillin-Streptomycin and maintained at 37 °C.

### Organoid Culture

Human PDAC organoids were generated in the Tuveson laboratory and cultured with assistance from the Organoids Shared Resource at Cold Spring Harbor Laboratory, as described (25). Briefly, organoids were passaged using H+++B Medium (Advanced DMEM/F-12 [Thermo Fisher], HEPES pH7.2-7.5 [10mM, Thermo Fisher], Glutamax [1x, Thermo Fisher], Primocin [100 μg/mL, Invivogen], BSA [0.1%, Sigma-Aldrich]. For organoids dissociation, cells were treated with TrypLE Express (Thermo Fisher) and then resuspend with Matrigel (Corning).

Human Complete Feeding Medium (hCPLT) (H+++B minus BSA, Wnt3a-conditioned medium [1x, cell line from ATCC], R-spondin1-conditioned medium [1x, cell line from Trevigen], B27 supplement [1x, Thermo Fisher], Nicotinamide [10 mM, Sigma-Aldrich], N-acetylcysteine [1.25 mM, Sigma-Aldrich], mNoggin [100 ng/mL, Peprotech], hEGF [50 ng/mL, Peprotech], hFGF [100 ng/mL, Peprotech], hGastrin I [10 nM, Tocris], A 83-01 [500 nM, Tocris] was added to organoids, and cultures were maintained at 37°C.

### ASO Delivery in 2D Cells

For ASO free-uptake in 2D cell culture, 2 × 10^4^ HEK293T cells were seeded in a 48-well plate and incubated overnight. On Day 1, the DMEM culture medium was changed and supplemented with the indicated ASO amounts. On Day 3, the medium was replaced with fresh medium plus ASO at the same final concentrations. On Day 5, the cells were harvested for RNA extraction. For ASO transfection, a single-cell suspension of HEK293T cells was prepared. Lipofectamine 3000 (Invitrogen) and ASO transfection mixes were prepared according to the manufacturer’s instructions in Opti-mem (Thermo Fisher). 2 × 10^4^ cells were incubated with transfection mix for 8 h in a 48-well tissue culture plate at 37 °C. After 8 h, the cells were transferred to a 15-mL conical tube and centrifuged for 5 min at 300 RCF at RT. The medium/Lipofectamine/ASO mixture was aspirated. The cells were then resuspended in fresh medium and plated in 48-well tissue-culture plates for 48 h at 37 °C.

### Free-uptake Delivery to Organoids

For ASO free-uptake in organoids, the organoids were passaged and prepared into single-cell suspensions using TrypLE. 5 × 10^4^ cells were resuspended in 40 μL of Matrigel and plated in a pre-warmed 24-well plate and incubated with hCPLT overnight. On Day 1, the medium was replaced with fresh medium supplemented with the desired concentration of ASO. On Day 3, the medium/ASO was replaced with fresh medium/ASO. On Day 5, the organoids were harvested for downstream assays.

### Free Uptake of ASO in Matrigel-Diluted Organoid Culture

Matrigel beds were made by combining equal volumes of DPBS (Thermo Fisher) and Matrigel in 96-well plates and incubating at 37 °C to form a Matrigel bed. Organoids were then dissociated into single cells and resuspended in culture medium with 10% Matrigel and ASOs at the indicated concentrations. 100 μL of organoid mixes (20 cells/μL) were plated on the Matrigel beds and incubated at 37 °C for 5 days. Organoids were harvested for downstream assays.

### ASO Transfection in Organoids

For ASO transfection in organoids, transfection mixes of ASO were prepared, and organoids were dissociated, as described above. 10^5^ cells/well were incubated in a 48-well plate_with Lipofectamine 3000 and ASO transfection mixes at 37 °C. After 8 h, the cells were transferred to 15-mL conical tubes and spun down. The supernatants were aspirated and the cell pellets were resuspended in Matrigel. The organoids were then plated and cultured at 37 °C for 72 h. Organoids were harvested for downstream assays.

### Western Blotting

Organoids were harvested and lysed in RIPA lysis buffer (50 mM Tris, pH 7.4, 150 mM NaCl, 0.1% SDS, 1% NP-40, 0.5% sodium deoxycholate and 1 mM PMSF + protease inhibitor cocktail (Roche)) by 30-min incubation on ice followed by sonicating for 3 min at medium power using a Bioruptor (Diagenode). Protein concentration was measured by Bradford assay (Bio-Rad) with BSA as a standard. Rabbit anti-PKM1 (1:500; Cell Signaling), rabbit anti-PKM2 (1:500; Cell Signaling), and mouse anti-Vinculin (1:1000, Santa Cruz Biotechnology) antibodies were used with IRDye 800CW secondary antibodies (LI-COR) for western blotting, and the blots were imaged and quantified using an Odyssey Infrared Imaging System (LI-COR).

### RNA Extraction, RT-PCR, and RT-qPCR

Total RNA was extracted with TRIzol (Life Technologies) according to the manufacturer’s protocol. Oligo dT(18)-primed reverse transcription was carried out with ImpromII Reverse Transcriptase (Promega). Semi-quantitative RT-PCR was carried out in the presence of [α-^32^P] dCTP (Perkin Elmer) with primer sequences provided in Supplementary Table 1. PCR products were digested with either DdeI or PstI for *SMN2* or *PKM2* detection, respectively. Digested products were then separated by 5% native PAGE, detected with a Typhoon FLA7000 phosphorimager, and quantified using ImageJ. Each band was normalized to the GC content and PSI was calculated.

For RT-qPCR, cDNA was analyzed on a QuantStudio 6 Flex Real-Time PCR system (Thermo Fisher). Fold changes were calculated using the ΔΔCq method. Primer sequences are listed in Supplemental Table 1.

### Pyruvate Kinase Assay

Organoids were transfected with ASO as described above. Organoids were pelleted and resuspended in PK Assay Buffer, and total protein concentration was determined via Bradford assay. Pyruvate Kinase assay was preformed according to the manufacturer’s instructions (Abcam). Optical density at 570 nm was measured at room temperature using a SpectraMax i3 plate reader (Molecular Devices) from 0 min to 1 h with 1-min interval time points.

## Figure Legends

**Figure S1:**
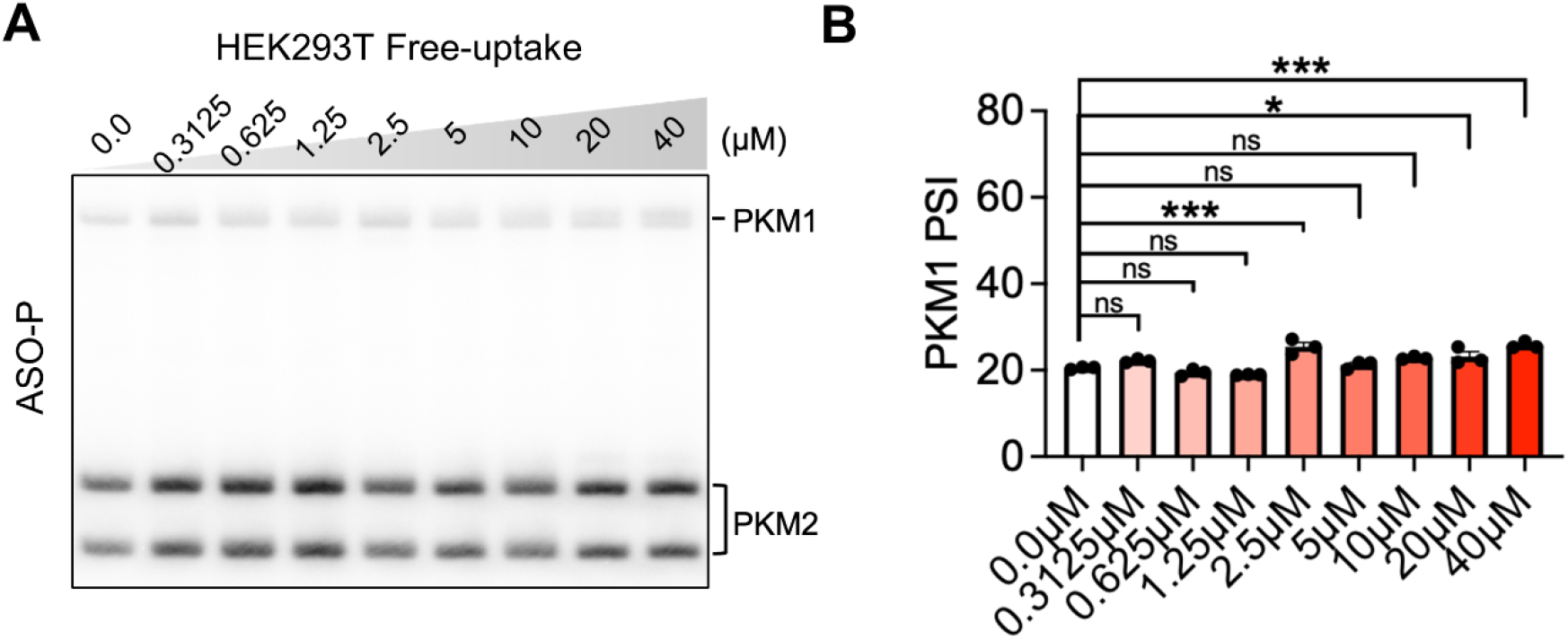
Free uptake of ASO-P in HEK293T Cells. **(A)** Radioactive RT-PCR of *PKM1/2* in HEK293T treated by free uptake of ASO-P. **(B)** PSI quantification of **(A)** (n=3). SC, scrambled control. One-way ANOVA, \**P*<0.05, \*\*\**P*<0.0001. Error bars represent ± SEM.

**Figure S2:**
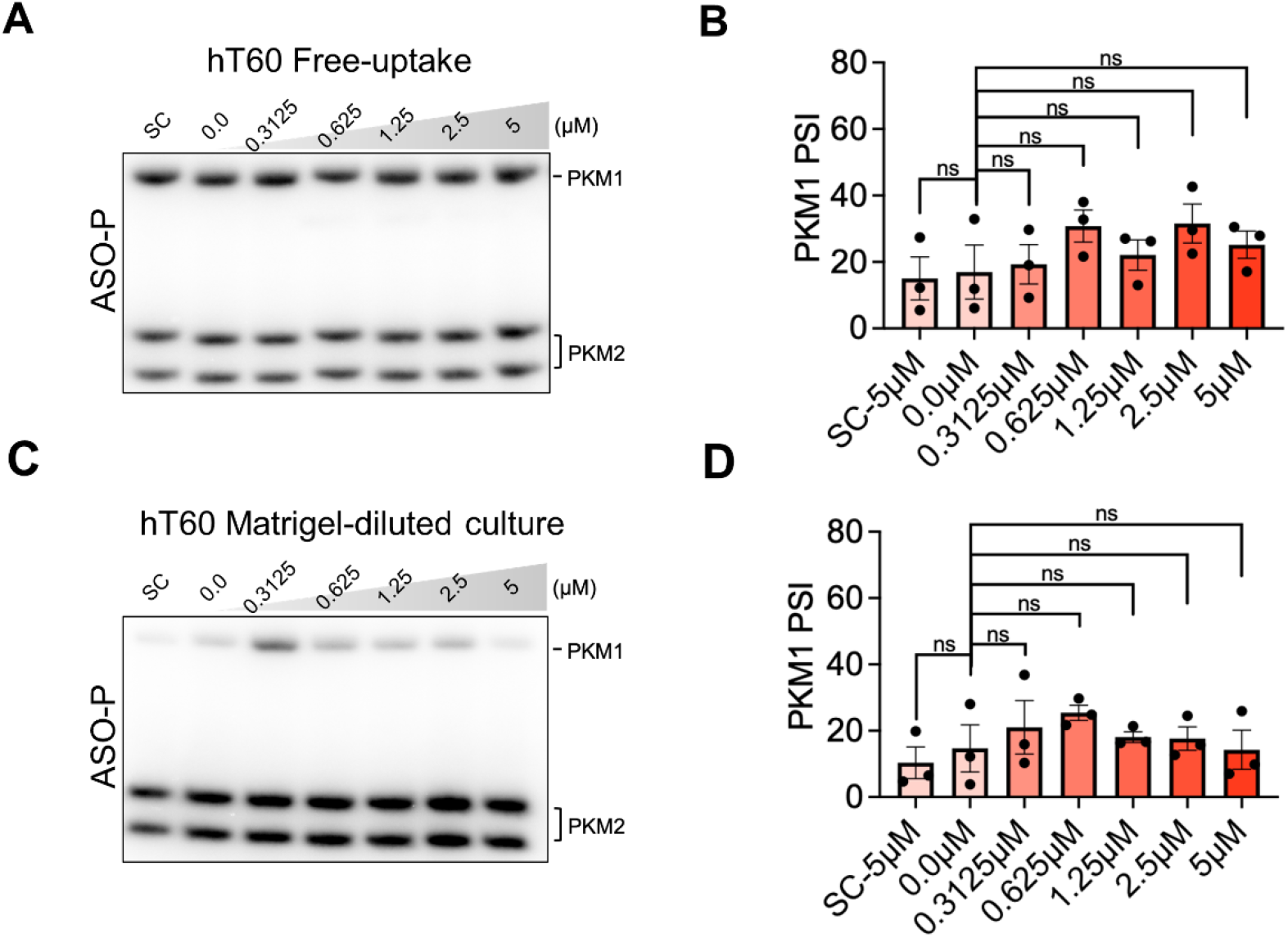
Free uptake of ASO-P in human hT60 organoid line. **(A)** Radioactive RT-PCR of *PKM1/2* mRNA in hT60 organoid treated by free uptake of ASO-P. **(B)** PSI quantification of **(A)** (n=3). **(C)** Radioactive RT-PCR of *PKM1/2* mRNA in Matrigel-diluted cultured hT60 organoids treated by free uptake of ASO-P. **(D)** PSI quantification of **(C)** (n=3). SC, scrambled control. One-way ANOVA. Error bars represent ± SEM.

**Figure S3:**
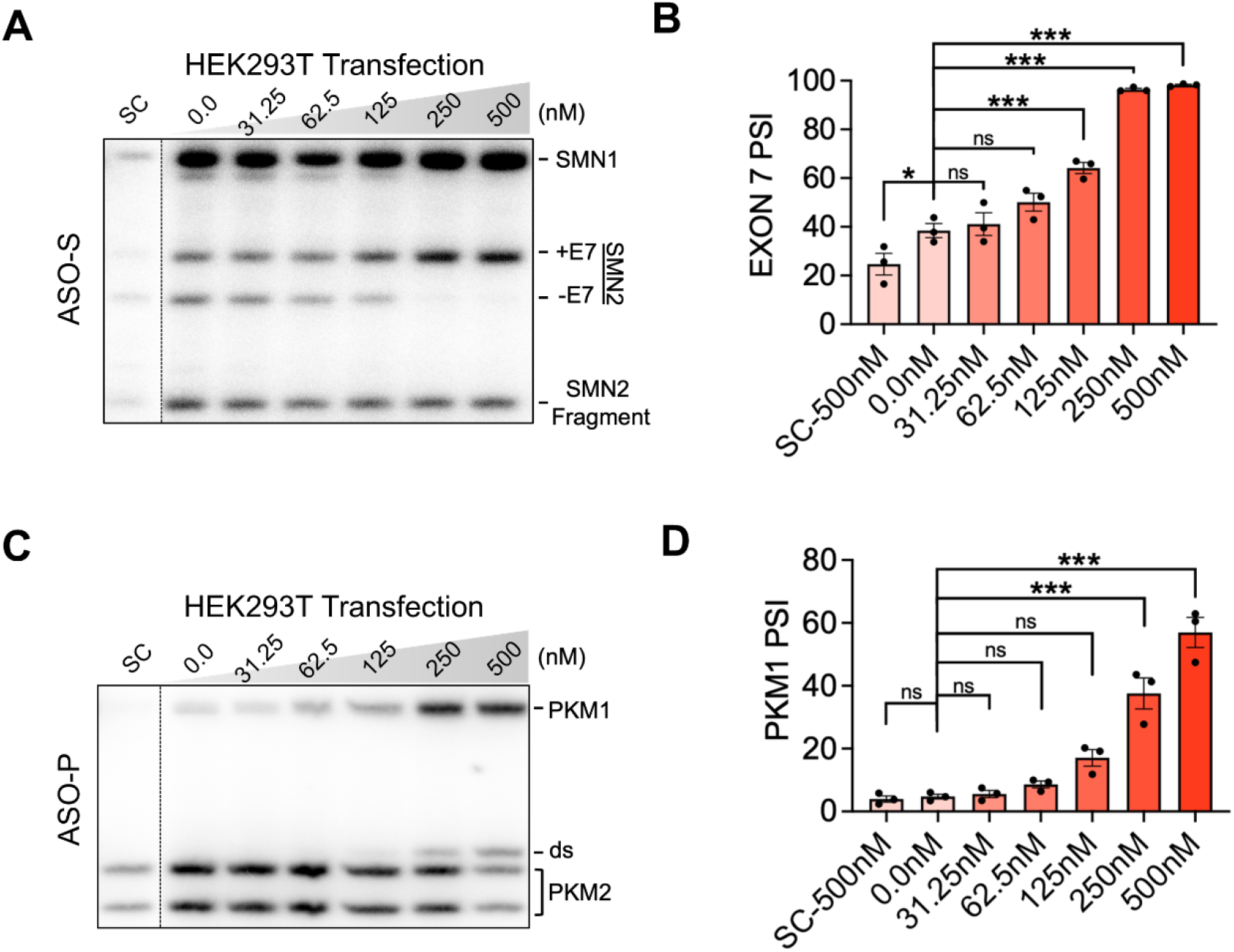
ASO transfection in human hT60 organoid line. **(A)** Radioactive RT-PCR of *PKM1/2* in hT60 organoid line transfected with ASO-P. **(B)** PSI quantification of **(A)** (n=3). **(C)** Western blot of PKM1, PKM2 in hT60 organoid line transfected with 125 nM ASO-P. Fold change was calculated by normalizing band intensity to vinculin (n=2). **(D)** Quantification of PK assay of hT60 organoid transfected with 125 nM ASO-P. PK activity and relative activity are shown (n=3). NTC, non-treatment control; SC, scrambled control. One-way ANOVA, \**P*<0.05, \*\*\**P*<0.001. Error bars represent ± SEM.

**Figure S4:**
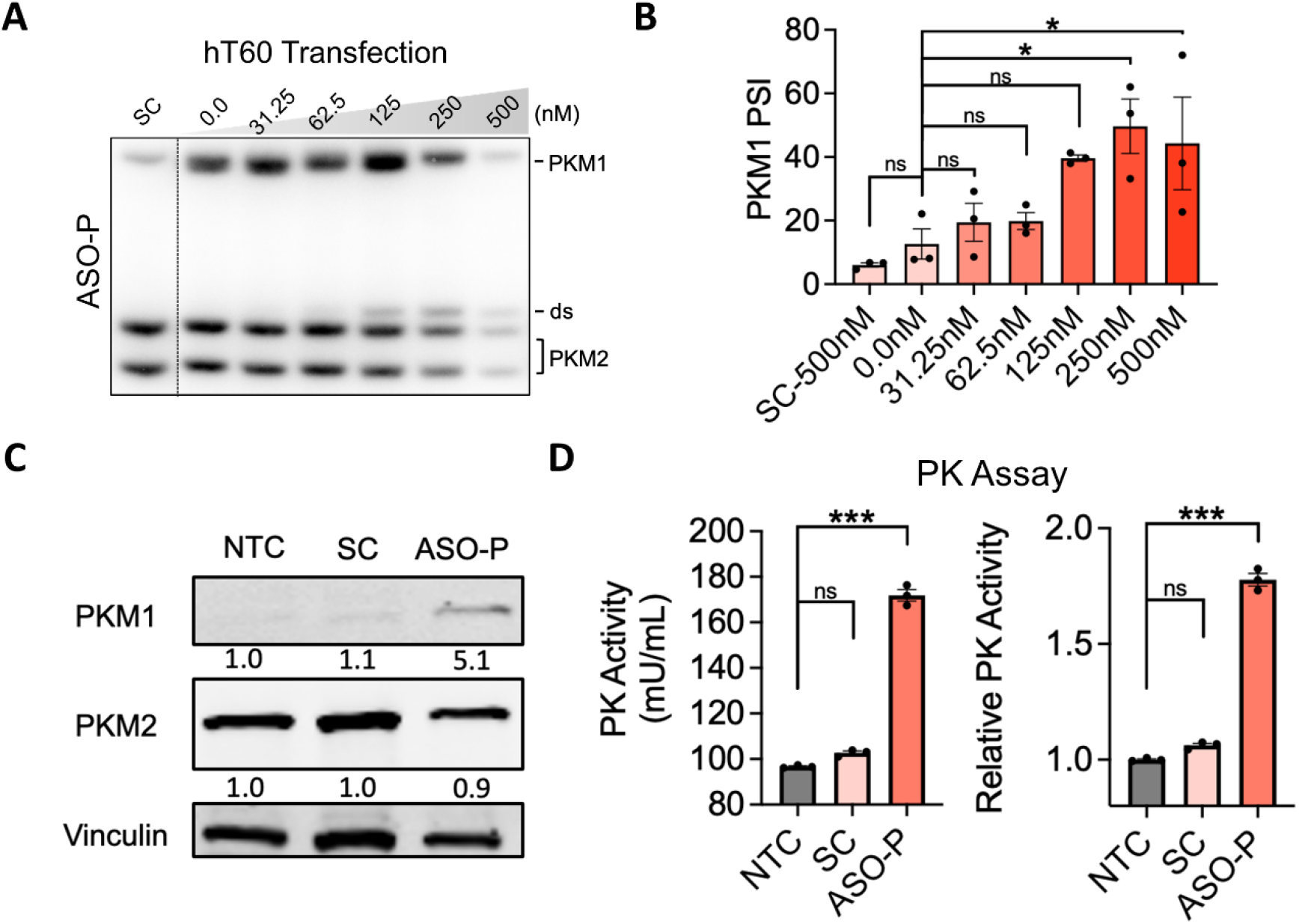
ASO transfection in HEK293T cells. **(A)** Radioactive RT-PCR of *SMN1/2* mRNA in HEK293T cells transfected with ASO-S. **(B)** PSI quantification of **(A)** (n=3). **(C)** Radioactive RT-PCR of *PKM1/2* mRNA in HEK293T cells transfected with ASO-P. **(D)** PSI quantification of **(C)** (n=3). SC, scrambled control. One-way ANOVA, \**P*<0.05, \*\*\**P*<0.0001. Error bars represent ± SEM.

---

**Table S1.**
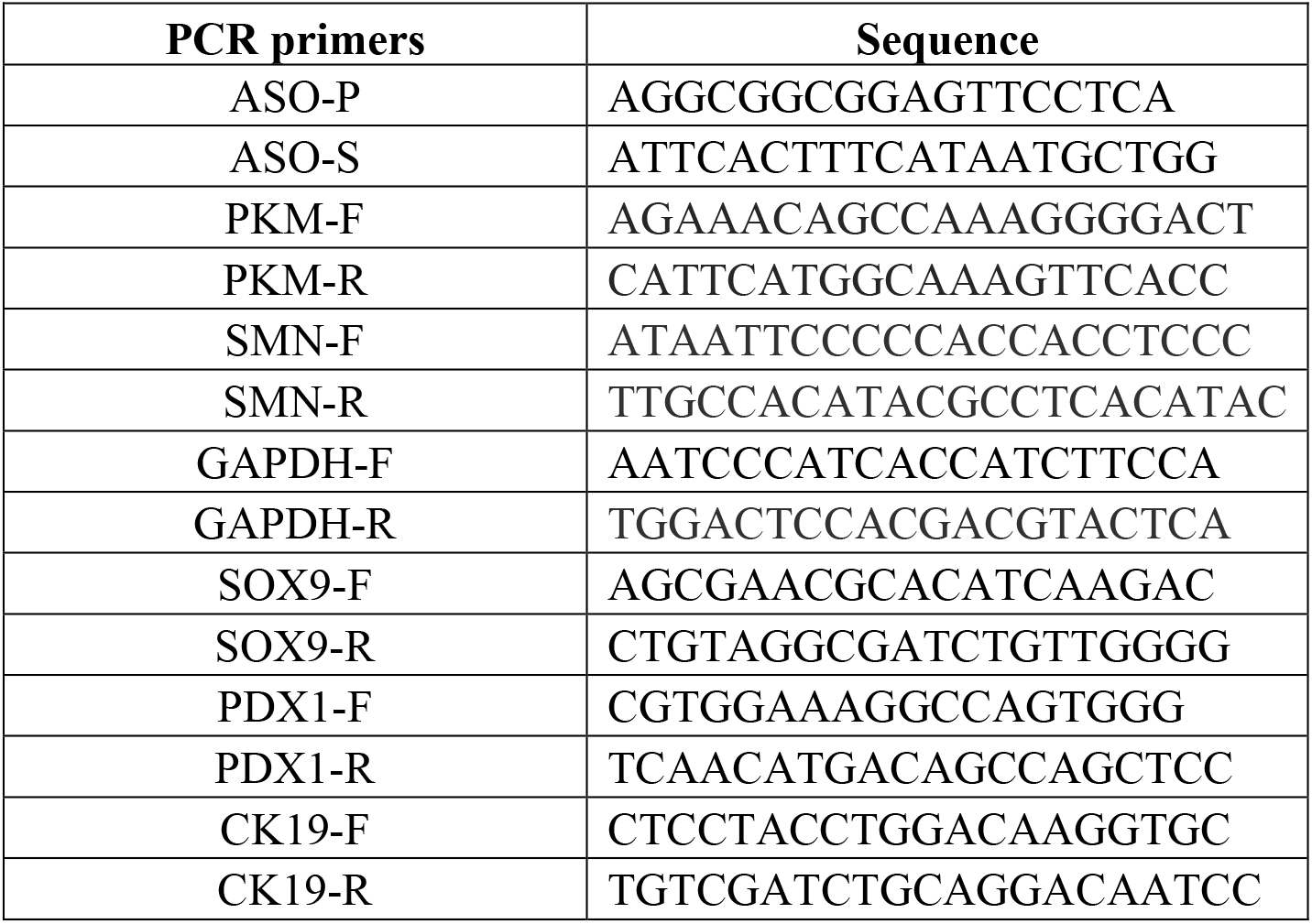
Oligonucleotides.

